# Validating wing biopsies for blood-borne pathogen characterization in bats

**DOI:** 10.64898/2026.03.11.711225

**Authors:** Molly C. Simonis, Amanda Vicente-Santos, Lauren R. Lock, Kristin E. Dyer, Beckett L. Olbrys, M. Brock Fenton, Karen E. Sears, Dmitriy V. Volokhov, Nancy B. Simmons, Daniel J. Becker

## Abstract

Wildlife surveillance is critical for tracking disease emergence, characterizing pathogen diversity, and assessing spillover risks. Blood-borne pathogens are of particular interest for such efforts due to their global distribution, broad host taxa, and zoonotic potential. Despite the need to monitor blood-borne pathogens, blood collection efforts are costly for both biologists and the wildlife being sampled (i.e., time-consuming and stressful), hindering our ability to expand and enhance surveillance efforts. There is thus a pressing need for reliable methods for detecting blood-borne pathogens that minimize sampling efforts and wildlife stress. Vascular tissues can contain enough blood to detect infections while minimizing sampling effort and stress on wildlife, but it is unclear how pathogen detection and characterization from these tissues compared to blood. To evaluate the reliability of using vascular tissues for detecting blood-borne pathogens in wildlife, we collected paired samples of blood and wing biopsies from individual common vampire bats (*Desmodus rotundus*) and molecularly screened them for bartonellae, hemotropic mycoplasmas (hemoplasmas), and trypanosomes. The probability of detection was consistently lower in wing tissues than in blood for all pathogens, possibly due to blood vessel avoidance when collecting the former. However, we detected infection in wing tissues of at least two individual bats for each blood-borne pathogen. Paired-positive individuals mostly showed high sequence concordance between tissues, indicating frequent detection of the same infections. Estimated sample sizes needed to detect a single infection and the reported prevalences were similar (i.e., *n* = 10–39). Due to the lower probability of infection in wing tissues compared to blood, we suggest that using these samples to estimate infection prevalence of blood-borne pathogens is not ideal. However, our results demonstrate that vascular tissues are viable for initial pathogen assessment and discovery to help target surveillance efforts in the future.

## Introduction

As emerging infectious diseases threaten both wildlife and human health [1,2], surveillance of wildlife pathogens is a critical aspect of tracking disease emergence, particularly zoonotic pathogens [3,4]. Characterizing wildlife pathogens for surveillance is important for managing and predicting disease outbreaks, epidemics, and pandemics [5], but there are many challenges associated with wildlife sampling for pathogen detection [6,7]. For example, efforts to survey wildlife can be costly in terms of time and efforts associated with animal capture (particularly for cryptic taxa), and invasive sampling strategies can cause additional stress to wildlife. Therefore, methods for field sampling and pathogen detection that reduce researcher effort and wildlife stress are important for sustainable, ongoing surveillance of zoonotic pathogens.

Blood-borne bacterial and protozoan pathogens are zoonoses of particular and growing concern [8–10], as they are geographically widespread across wildlife taxa [11–19]. Bacterial bartonellae and hemotropic mycoplasmas (hemoplasmas), as well as protozoan trypanosomes, have varying routes of transmission. For example, bartonellae in bats have evidence of being largely vectored by arthropods [20], but *Bartonella henselae*, which causes cat scratch fever, is typically directly transmitted to humans through cat bites and scratches (although vectored transmission is still possible [21]). Hemoplasmas show evidence of direct transmission between bats and humans (via bat consumption [22,23]), vertical transmission within small mammalian host species [9], and potential vector transmission [24]. Trypanosomes are commonly vector-transmitted via triatomines and flies as well as through direct oral transmission [25,26]. Due to varying transmission routes, hosts, and evidence of cross-species transmission, we need surveillance of these blood-borne bacterial and protozoan pathogens to better understand their genetic diversity in wildlife and spillover risks. Surveillance of blood-borne pathogens in wildlife ideally involves detecting infections from blood samples, but collecting blood from wildlife can be time-consuming, requires specialized training, and can place significant stress on individuals being sampled [27–29]. Blood-borne bacterial and trypanosome infections can be detected in functional organ tissues such as the spleen, liver, and heart [30–32]. However, collecting organs can be time-intensive and requires lethal sampling, which may run counter to conservation concerns, conflicts with other ongoing wildlife studies, and permitting issues.

Vascular tissue sampling methods for blood-borne pathogens that are logistically easier, less time-consuming, and non-lethal, such as earclips from domestic cats and cattle [33,34], ear biopsies and toe clips in rodents [35,36], or wing tissues in bats [37–39], may contain enough blood to detect infections and also minimize sampling effort and stress on individual wildlife. Unlike blood samples, tissue samples such as these are routinely collected and vouchered in museums [40,41], many of which are increasingly stored cryogenically [42], so tissues may be accessible without sampling new individuals in the field. However, it is unclear how low-effort vascular tissue sampling compares with direct blood collection for our ability to detect and identify blood-borne pathogens.

Bats are commonly associated with emerging zoonoses, including blood-borne pathogens such as bartonellae, hemoplasmas, and trypanosomes [12,22,23,38,43,44]. Bat wings are highly vascularized, and wing tissues are commonly collected for molecular species identification [45], stable isotope analyses [46], characterizing population structure [47], quantifying aging [48–50], and -omics studies of species diversity [51]. While bat wing tissues have been used to detect blood-borne pathogens previously [37,39], it is unclear how using wing tissues instead of blood could facilitate surveillance.

Here, we use paired blood and wing tissues from common vampire bats (*Desmodus rotundus*) collected in Belize to determine concordant infection detection across the two sample types. We used tissues from each individual to investigate detection probability for infections with bartonellae, hemoplasmas, and trypanosomes, all of which are known to occur with high prevalence in the local vampire bat population [13,52–56]. We hypothesized that the probability of detecting infection would be lower in wing tissue samples compared to blood tissues due to collection methods (i.e., researchers typically avoid visible wing blood vessels when collecting biopsies) and blood volumes in those samples compared to whole blood.

## Methods

### Vampire bat sampling

During two, two-week trips in November 2021 and April–May 2022, we sampled common vampire bats in Orange Walk District, Belize as part of a long-term study of the local bat community [13,57,58]. Bats were captured using mist nets and harp traps along flight paths within the Lamanai Archeological Reserve and Ka’Kabish Archaeological Site. Bats were marked with 3.5 mm incoloy wing bands (Porzana Inc.), and we obtained blood by lancing the propatagial vein with 23-gauge needles, followed by collection in heparinized capillary tubes. Blood was stored on Whatman FTA cards and at ambient temperature in the field prior to long-term preservation at −20 °C at the University of Oklahoma. We also collected two 2 mm wing biopsy punches along the lower wing membrane close to the body wall and stored them in 1 mL DNA/RNA Shield at −80 °C. All bats were released following sampling.

Field procedures were performed according to guidelines for the safe and humane handling of bats published by the American Society of Mammalogists [59] and were approved by the Institutional Animal Care and Use Committees of the American Museum of Natural History (AMNH IACUC-20210614) and the University of Oklahoma (2022–0197). All bat sampling was authorized by the Belize Forest Department through scientific collection permit FD/WL/1/21(12) and Belize Institute of Archaeology permits IA/S/5/6/2l(01) and IA/H/1/22(03).

### DNA extraction

As described previously, we extracted genomic DNA from blood on Whatman FTA cards using QIAamp DNA Investigator Kits (Qiagen) [13,57,58]. From a broader collection of blood samples, we here focused analyses on individual bats with paired blood and wing tissues (*n* = 63). Sample availability enabled screening 60 individuals for infections with bartonellae and trypanosomes, while 62 individuals were screened for bartonellae, trypanosomes, and hemoplasmas. Due to ongoing projects across institutions using wing tissues, we used two different DNA extraction methods. Most wing tissues (*n* = 45) were processed at University of Oklahoma using the *Quick*-DNA/RNA Viral Magbead Kit (Zymo Research) run on an IsoPure 96 platform (Accuris Instruments). Briefly, we used a vortex mixer to homogenize wing tissues and included a 15-minute proteinase K digestion at room temperature per the manufacturer’s instructions. DNA was eluted into 50 µL nuclease-free water. The remaining wing tissues (*n* = 18) were processed at University of California Los Angeles using the *Quick*-DNA Miniprep Plus Kit (Zymo Research), following the manufacturer’s solid tissue protocol. Prior to extraction, we used a rotor-stator homogenizer to homogenize wing tissues followed by an overnight proteinase K digestion at 55°C. DNA was eluted into 30 µL nuclease-free water. DNA extraction methods for wing tissues yielded 0.02 ng/μL greater concentrations when using the *Quick*-DNA Miniprep Plus Kit compared to the *Quick*-DNA/RNA Viral Magbead Kit (generalized linear model [GLM] with Tweedie response: F_1_ = 40.1, P < 0.0001, R^2^ = 0.47). However, DNA purity (260/280) did not differ between extraction methods (GLM with Tweedie response: F_1_ = 0.62, P = 0.43, R^2^ = 0). Finally, DNA concentrations were 0.11 ng/μL greater in blood than wing tissues when analyzing paired samples (generalized linear mixed model [GLMM] with Tweedie response and random effect of individual bat: F_1, 63_ = 12.94, P < 0.001; R^2^ = 0.40), but DNA purity did not differ between paired samples (GLMM with the same structure: F_1, 59_ = 2.1, P = 0.15, R^2^ = 0.57).

### Pathogen screening

We used previously established protocols to screen all paired blood and wing tissue DNA samples for bartonellae, hemoplasmas, and trypanosomes (Table S1). For bartonellae and trypanosomes, we used nested PCR to target the *gltA* and small subunit rRNA (SSU rRNA) genes, respectively [60–62]. While the former PCR is highly specific to bartonellae, the latter PCR amplifies all trypanosomatids except salivary trypanosomes, unlike methods targeting *Trypanosoma cruzi* specifically [63]. Amplicons were purified with Zymo kits (DNA Clean and Concentrator-5, Zymoclean Gel DNA Recovery) and directly Sanger sequenced at the North Carolina State University Genomic Sciences Laboratory. For hemoplasmas, we used PCR targeting the 16S rRNA gene [12,13,53], with amplicons purified and directly Sanger sequenced at Psomagen. Forward and reverse sequences were trimmed and cleaned in Geneious [64], and we used NCBI BLASTn and previous protocols to assign consensus sequences to genotypes [12,52,65,66]. Data for bartonellae and hemoplasmas in blood have been published previously [52], whereas trypanosome data for blood and data for all pathogens in wing tissues were generated here. Infection positivity in wing tissues did not differ by extraction methods for bartonellae, but was greater for hemoplasmas using wing tissues extracted via the *Quick*-DNA/RNA Viral Magbead Kit compared to the *Quick*-DNA Miniprep Plus Kit (GLM with Tweedie response for each pathogen; Table S2). There was also an effect of extraction method on trypanosome infection positivity, but a post hoc pairwise contrast indicated no difference (GLM with Tweedie response; Table S2). As such, we controlled for these differences in our subsequent analyses of paired blood and wing tissues (see *Statistical analyses* below).

### Statistical analyses

All statistical analyses and data visualizations were performed in the statistical environment R (version 4.3.1, Beagle Scouts) and the package *ggplot2* [67,68]. To determine how infection detection differed between blood and wing tissues, we fit binomial GLMMs using the package *lme4* [69] for infection status of each pathogen as a function of sample type and a random effect of individual bat to account for multiple measures (each individual had a blood and wing tissues sampled). Given the above differences in pathogen detection for wing tissues extracted using different kits, we also included a random effect of extraction method in our analyses [70]. We tested each model via a type-II ANOVA from the *car* package [71], and we quantified post-hoc estimated marginal means and confidence intervals using the *emmeans* package [72]. We present all GLMM coefficients exponentiated as odds ratios (ORs).

Finally, to determine the sample size needed to detect infections in wing tissues, we calculated binomial probabilities using a 95% confidence level and 80% statistical power. We first estimated the minimum sample size for detecting a single infection in wing tissues, such that the probability of zero detections was denoted by (1 − *p*)^*n*^, and the probability of a single detection was denoted by 1 − (1 − *p*)^*n*^, where *p* is the resulting infection probability of bartonella, hemoplasma, or trypanosome infections found here. We then also estimated the sample size needed to detect the observed prevalence of bacterial and protozoan pathogens found here using *pwr* package in R [73], again assuming a 95% confidence level in detection probability and 80% statistical power.

## Results

Of the 63 bats screened (60 for bartonellae and trypanosomes, 62 for hemoplasmas), 49 were positive for baronellae (mean [lower CI, upper CI]; 82% [70%, 89%]); 48 positive from blood, three positive from wing tissues), 38 were positive for hemoplasmas (61% [49%, 72%]; 33 positive from blood, nine positive from wing tissues), and 26 were positive for trypanosomes (43% [32%, 56%]; 24 positive from blood, two positive from wing tissues). One, four, and one bat were positive in wing tissues but not blood for bartonellae, hemoplasmas, and trypanosomes, respectively. Fourteen bats were coinfected with all three pathogens, two of which would not have been detected without wing tissue screening. One individual was coinfected with bartonellae and trypanosomes only. Two individuals were positive for bartonellae in both blood and wing tissues, and five were positive for hemoplasmas in both tissues (Table 1). For individuals that tested positive in both blood and wing tissue, bartonellae sequences were 100% similar between blood and wing tissues, whereas hemoplasma sequences were 98.66% ± 0.54% (mean ± SE) similar between samples (Table 1).

**Table 1.** Percent similarity between genetic pathogen sequences identified in all individual vampire bats with paired infection positivity (i.e., infections found in both blood and wing tissues). BZID represents the unique identifier for each individual bat. Lineage match indicates if paired infections had the same genotype. Note: While the study detected trypanosome infections in wing tissues from two individuals, no cases showed paired positivity (infection detected in both blood and wing tissues from the same individual), therefore no trypanosome sequences are included in this genetic concordance analysis.

| BZID | Pathogen | Blood accession and identity | Wing accession and identity | Percent similarity* | Lineage match |
| --- | --- | --- | --- | --- | --- |
| 123 | bartonellae | PQ758887; DR11 genotype | PV599394; DR11 genotype | 100 | Y |
| 1435 | bartonellae | PQ758926; DR1 genotype | PV599395; DR1 genotype | 100 | Y |
| 138 | hemoplasma | OQ546566; VB2 genotype | PX705794; VB2 genotype | 99.7 | Y |
| 1598 | hemoplasma | OQ385172; VB1 genotype | PX705791; VB2 genotype | 97.2 | N |
| 372 | hemoplasma | OQ546522; VB1 genotype | PX705793; VB1 genotype | 99.8 | Y |
| 79 | hemoplasma | OQ546567; VB3 genotype | PX705797; VB3 genotype | 99.0 | Y |
| 1042 | hemoplasma | OQ385170; VB1 genotype | PX705792; VB2 genotype | 97.6 | N |

\*Genetic similarity between blood and wing tissue sequences

The probability of detecting infection was consistently lower in wing tissues than in blood samples across all three pathogens. However, despite lower detection probabilities for wing tissues compared to blood, we did detect at least two positives in wing tissue for each pathogen. Detecting infections of bartonellae in wing tissues (0.05 [0.02, 0.14]) were 98% less likely than in blood (0.77 [0.65, 0.86]; OR = 0.02, P < 0.0001, R^2^_m_ = 0.57, R^2^_c_ = 0.57). Detecting hemoplasma infections in wing tissues (0.15 [0.05, 0.38]) were 92% less likely than in blood (0.69 [0.38, 0.89]; OR = 0.08; P < 0.0001, R^2^_m_ = 0.28, R^2^_c_= 0.41). Finally, detecting trypanosome infections in wing tissues (0.03 [0.01, 0.12]) were 95% less likely than those in blood (0.39 [0.27, 0.51]; OR = 0.05, P = 0.0001, R^2^_m_ = 0.40, R^2^_c_ = 0.40; Figure 1).

**Figure 1.**
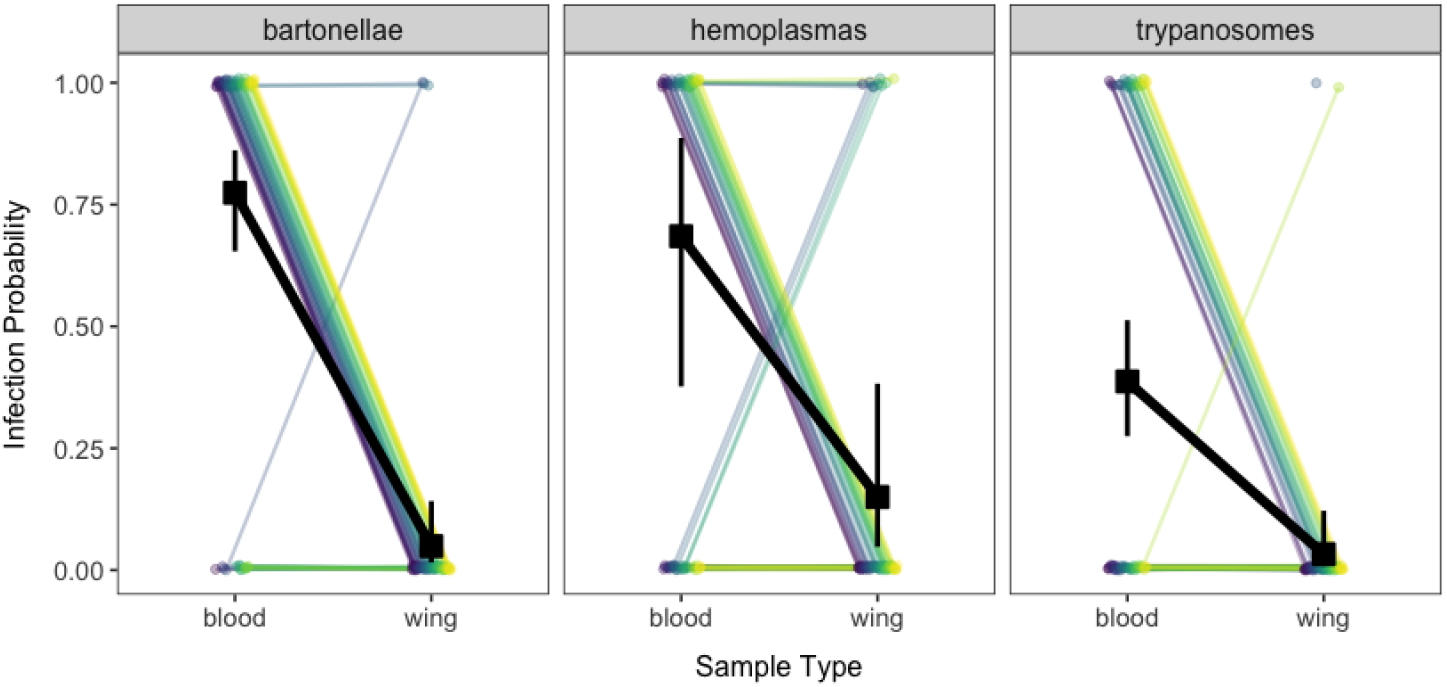
Probability of infection between blood and wing tissues in vampire bats when screened for bartonellae (left; n = 60), hemoplasmas (middle; n = 62), and trypanosomes (right; n = 60). Colored points and lines represent paired detection results for each individual bat. Black points, bars, and lines represent mean infection probabilities and 95% confidence intervals from the respective GLMMs across sample types.

The minimum estimated sample size to detect a single infection in wing tissues with 80% statistical power and 95% confidence was 31, 10, and 32 for bartonellae, hemoplasmas, and trypanosomes, respectively (Figure 2). The minimum estimated sample size to obtain an expected prevalence of 5% (bartonellae), 15% (hemoplasmas), or 5% (trypanosomes) with 80% statistical power and 95% confidence was 39, 13, and 39, respectively (Figure 2).

**Figure 2.**
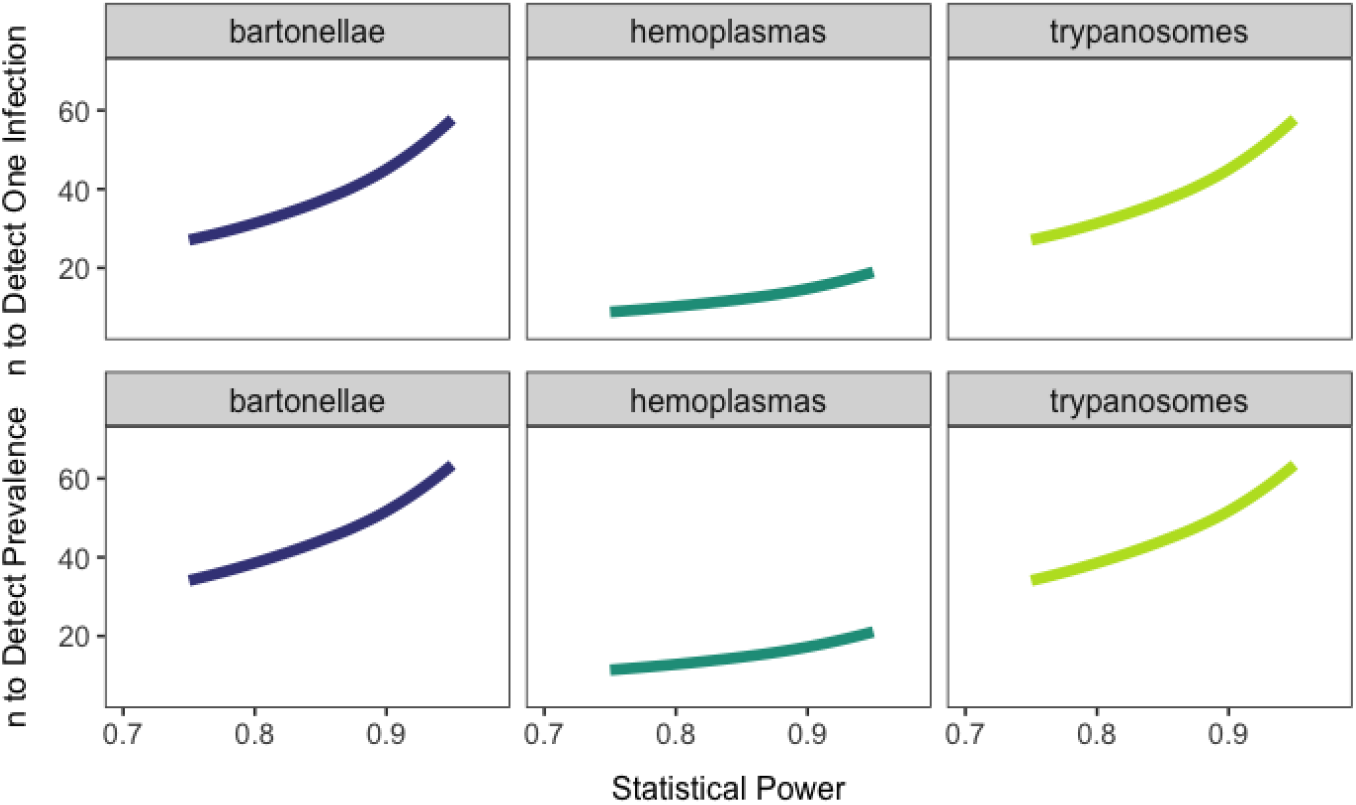
Estimated sample sizes needed to detect a single infection (top) and observed prevalence (bottom) for bartonellae, hemoplasmas, and trypanosomes detected in wing tissues.

## Discussion

Pathogen discovery and surveillance are critical for predicting and managing emerging infectious diseases. Capturing and sampling wildlife can be time and labor intensive as well as stressful for study animals, so identifying field methods that minimize researcher sampling effort and wildlife stress is key to efficient and humane pathogen monitoring. Here, we detected blood-borne pathogens in vampire bats using paired blood and wing tissue samples. Our results demonstrate that wing tissues represent a biologically relevant sample type for pathogens that requires less researcher effort and physiological stress to bats than does blood collection.

The odds of detecting blood-borne pathogens in wing tissues were substantially lower than blood samples, with the likelihood of detecting these infections being 98%, 92%, and 95% less likely for bartonellae, hemoplasmas, and trypanosomes, respectively. While we expected this reduced detectability given that researchers typically avoid blood vessels when collecting wing tissues in bats, we show here that wing tissues are still a viable option for initial pathogen detection and genetic characterization. Our detection probabilities are comparable to those reported in other studies that used wing tissues for detecting blood-borne pathogens. For example, trypanosomes from wing tissues were detected in four of 17 *Desmodus rotundus* in Brazil [30] and one of 361 wing tissues from *Tadarida brasiliensis* (Mexican free-tailed bats) in Oklahoma, USA [37], whereas we detected trypanosomes in three of 60 wing tissues. While we found that infection probabilities were consistently higher in blood samples than in wing tissues, all three pathogens were detectable to some degree in this sample type. DNA sequences from pathogens identified in wing tissues typically matched those from blood in the same individual bat, indicating the same infections were detected. We therefore suggest that wing tissues could be suitable for initial pathogen screening assessments and help guide further surveillance efforts. Despite the lower prevalence in wing tissues, such tissues are especially promising for blood-borne pathogen surveys of bats, given that they are easy to collect in large numbers and are commonly collected in field surveys.

Using bat wing tissues for detecting blood-borne pathogens could be applied more widely in pathogen discovery efforts. Given that wing tissues are often collected for use in molecular species identification [45], stable isotope analyses [46], characterizing population structure [47], quantifying aging [48–50], and -omics studies of species diversity [51], there are a vast number of wing tissue samples stored across the bat research community. Similarly, wing tissues are commonly collected during routine sampling expeditions for museum collections. Future studies could compare infection detection in wing tissues across multiple storage methods (e.g., ethanol, formalin-fixed, silica beads, DNA/RNA Shield) to ensure successful application for pathogen discovery more broadly. These samples could be used for initial assessments to determine which blood-borne pathogens may be circulating. Researchers could then decide whether additional efforts are needed for targeted surveillance (e.g., longitudinal study), using blood collection to obtain more accurate estimates of infection prevalence and better characterize genetic variants and determine zoonotic risks.

We used two different but methodologically similar approaches to extract DNA from wing tissues. DNA concentrations were, on average, 0.11 ng/μL greater when extracted using the *Quick*-DNA Miniprep Plus Kit compared to the *Quick*-DNA/RNA Viral Magbead Kit. We suggest this difference in concentrations is negligible, given that DNA purity did not differ between extraction methods and infection positivity did not differ between methods for both bartonellae and trypanosome infections. Although wing tissues extracted using the *Quick*-DNA/RNA Viral Magbead Kit were 86% more likely to be positive for hemoplasmas than those extracted using the *Quick*-DNA Miniprep Plus Kit, we suggest this difference is more likely due to the sample size imbalance between kits used and the general rarity of hemoplasma infections in wing tissues. Therefore, blood-borne infection testing from wing tissues are likely reliable regardless of extraction methods used.

We were able to detect infections of bartonellae, hemoplasmas, and trypanosomes in at least two vampire bats for each pathogen. Six infections would not have been detected without wing tissue screening (one infection with bartonellae, four hemoplasma infections, and one trypanosome infection). However, our ability to detect infections in wing tissue varied by pathogen (i.e., three infections with bartonellae, nine hemoplasma infections, and two trypanosome infections), and achievement of genetic concordance in individuals with paired blood and wing infections was limited to bartonellae only. Limitations in wing tissue detections and genetic concordance could be due to the tropism spectrum of each pathogen. For example, bartonellae typically infect red blood cells and endothelial walls of blood vessels [74,75], and hemoplasmas are intracellular infections of red blood cells (although they can also infect reproductive tissues) [76]. Trypanosomes are known for varied tissue tropism across organ systems in mammalian hosts [77]. Taken together, the specificity of tropism for bartonellae and hemoplasmas in blood and blood-related tissue may have contributed to our ability to detect more infections in vascular wing tissue compared to trypanosome infections. These limitations in infection detections suggest, again, that using wing tissues for pathogen detection may be best suited for pathogen discovery instead of general surveillance and estimating infection prevalence.

While blood-borne pathogen detection in wing tissues may be limited, we quantified the sample sizes needed to obtain bartonellae, hemoplasma, or trypanosome infections for future pathogen discovery. Sample sizes needed to obtain a single infection and observed prevalence here were similar. With wing tissues in bats being collected and used for a variety of bat research topics, future collaborative sampling should contribute to better understanding the genetic diversity of these geographically widespread pathogens.

In conclusion, we show that using bat wing tissues for initial pathogen assessment and discovery is a viable and less-invasive sampling alternative to bat blood. Although detecting blood-borne pathogens in wing tissues is likely not useful for estimating infection prevalence, sampling wild bat wing tissues for initial pathogen assessments and discovery can help guide more targeted and longitudinal surveillance efforts. We recommend bat biologists begin using wing tissues for blood-borne pathogen detection to expand our fundamental knowledge of wildlife pathogen diversity and spillover risks.

## Acknowledgements

We thank the late Mark Howells and the staff of the Lamanai Field Research Center for assistance with field logistics and permits as well as the many colleagues who helped net bats during 2021 and 2022 research in Belize.

## Funding

This study was supported by the Explorers Club, National Institutes of Health (R03AI188200), Edward Mallinckrodt, Jr. Foundation, National Science Foundation (DBI 2515340, DEB 2508535), and Research Corporation for Science Advancement (RCSA), Subaward No. 28365, part of a USDA Non-Assistance Cooperative Agreement with RCSA Federal Award No. 58-3022-0-005. MCS was supported by an appointment to the Intelligence Community Postdoctoral Research Fellowship Program at University of Oklahoma administered by Oak Ridge Institute for Science and Education through an interagency agreement between the U.S. Department of Energy and the Office of the Director of National Intelligence.

## Data availability

Sequence data are available on GenBank under accessions PX794553, PX794555, PX794557-PX794576 (trypanosomes in blood); PV599394–PV599396 (bartonellae in wing tissues); PX705789-PX705797 (hemoplasmas in wing tissues); and PX794554 and PX794556 (trypanosomes in wing tissues). Sequence data for bartonellae and hemoplasmas in blood have been provided previously [58]. Positivity data and all R code are available via Zenodo [78].

## Notes

https://doi.org/10.5281/zenodo.18947639

### Competing Interest Statement

The authors have declared no competing interest.

