## Supporting Information for "Validating wing biopsies for blood-borne pathogen characterization in bats"

**Running head:** Bat tissue concordance for pathogen surveillance

**Keywords:** bartonellae, hemoplasmas, trypanosomes, vampire bat

### Supporting Information

**Table S1.** PCR protocols and primers used for infection detection in both blood and wing tissues.

| PCR Protocol | | Primers | Reaction | Thermocycler Settings | References |
|---|---|---|---|---|---|
| bartonellae (*gltA* gene); nested | PCR1 | Forward: 443f<br>5'-GCTATGTCTGCATTCTATCA-3'<br>Reverse: 1210r<br>5'-GATCYTCAATCATTTCTTTCCA-3' | 12.5 µL Promega GoTaq® Green<br>1 µL 443f<br>1 µL 1210r<br>7.5 µL Nuclease Free H2O<br>3 µL DNA template<br>(25 µL total volume) | 2 min at 95 °C, 30 sec at 95 °C, 30 sec at 48 °C, 2 min at 72 °C (40 cycles), 7 min at 72 °C | (Bai et al., 2016; Becker et al., 2021) |
| | PCR2 | Forward: 781f<br>5'-GGGGACCAGCTCATGGTGG-3'<br>Reverse: 1137r<br>5'-AATGCAAAAAGAACAGTAAACA-3' | 12.5 µL Promega GoTaq® Green<br>1 µL 781f<br>1 µL 1137r<br>9.5 µL Nuclease Free H2O<br>1 µL PCR1 Product<br>(25 µL total volume) | 3 min at 95 °C, 30 sec at 95 °C, 30 sec at 55 °C, 2 min at 72 °C (40 cycles), 7 min at 72 °C, Read at ~300 base pairs | |
| trypanosomes (pan-trypanosome); nested | PCR1 | Forward: TRY927F<br>5'-CAGAAACGAAACACGGGAG-3'<br>Reverse: TRY927R<br>5'-CCTACTGGGCAGCTTGGA-3' | 12.5 µL Promega GoTaq® Green<br>0.5 µL TRY927F<br>0.5 µL TRY927R<br>6.5 µL Nuclease Free H2O<br>5 µL DNA template<br>(25 µL total volume) | 5 min at 95 °C, 30 sec at 94 °C, 30 sec at 55 °C, 1 min at 72 °C (33 cycles), 10 min at 72 °C | (Noyes et al., 1999) |
| | PCR2 | Forward: SSU561F<br>5'-TGGGATAACAAAGGAGCA-3'<br>Reverse: SSU561R<br>5'-CTGAGACTGTAACCTCAAAGC-3' | 12.5 µL Promega GoTaq® Green<br>0.5 µL SSU561F<br>0.5 µL SSU561R<br>9.5 µL Nuclease Free H2O<br>2 µL PCR1 Product<br>(25 µL total volume) | 5 min at 95 °C, 30 sec at 94 °C, 30 sec at 55 °C, 45 sec at 72 °C (33 cycles), 10 min at 72 °C, Read at ~650 base pairs | |

| | | | | | |
|---|---|---|---|---|---|
| hemoplasmas (16S rRNA gene) | | Forward: UNI_16S_hemoFnew 5'-TGAATAAGTGACAGCWAACTATGTGCC-3' Reverse: UNI_16S_hemoR 5'-GACGGGCGGTGTGTACAAGACCTG-3' | 24.5 µL Promega GoTaq® Green 3.5 µL UNI_16S_hemoFnew 3.5 µL UNI_16S_hemoR 8.5 µL Nuclease Free H2O 10 µL DNA template (50 µL total volume) | 5 min at 95 °C, 50 sec at 96 °C, 1 min at 60 °C, 1 min at 72 °C (55 cycles), 7 min at 72 °C, Read at ~850-900 base pairs | (Becker et al., 2025; Volokhov et al., 2017) |

**Table S2.** Generalized linear models and outcomes for determining differences in infection positivity between wing tissue extraction methods using *Quick*-DNA/RNA Viral Magbead Kit (control) compared to the *Quick*-DNA Miniprep Plus Kit. All models were fit using a binomial response.

| Model | n | df | odds ratio | P value (main effect) | P value (contrast) | $R^2$ |
|---|---|---|---|---|---|---|
| wing bartonellae infection status ~ extraction method | 61 | 1 | 0.76 | 0.83 | 0.83 | 0.005 |
| wing hemoplasma infection status ~ extraction method | 63 | 1 | 0.14 | 0.01 | 0.01 | 0.19 |
| wing trypanosome infection status ~ extraction method | 61 | 1 | < 0.0001 | 0.02 | 1.00 | 0.96 |
